# Antigen-Specific T Cell Receptor Discovery for Treating Progressive Multifocal Leukoencephalopathy

**DOI:** 10.1101/2024.11.04.621904

**Authors:** Sasha Gupta, Tijana Martinov, Ashley Thelen, Megumi Sunahara, Shwetha Mureli, Angie Vazquez, Josiah Gerdts, Ravi Dandekar, Irene Cortese, Camille Fouassier, Elaine Schanzer, Fyodor D. Urnov, Alexander Marson, Brian R. Shy, Philip D. Greenberg, Michael R. Wilson

**Author notes:** **Relevant Disclosures, *if author not listed then none reportable*:** T.M. is an inventor on patents with rights held by the Fred Hutchinson Cancer Center pertaining to TCR discovery; two of the patents are licensed to Affini-T Therapeutics. T.M. and A.T. are currently employees of Bristol Myer Squib. I.C. is shareholder for Nouscom, AG; Keires, AG. B.R.S is a member of the scientific advisory board for Kano Therapeutics. B.R.S. is an inventor on patents with rights held by the Regents of the University of California pertaining to genome editing in primary hematopoietic cells. FDU is a paid consultant to Vertex Pharmaceuticals, to Ionis Pharmaceuticals, receives salary and research support from Danaher Corporation, owns equity in and is a compensated advisor to Tune Therapeutics and Cimeio Therapeutics, and is a named inventor on a number of patent filings in the CRISPR-Cas therapeutic space that are owned by the Regents of the University of California. A.M. is a cofounder of Site Tx, Arsenal Biosciences, Spotlight Therapeutics and Survey Genomics, serves on the boards of directors at Site Tx, Spotlight Therapeutics and Survey Genomics, is a member of the scientific advisory boards of Site Tx, Arsenal Biosciences, Cellanome, Spotlight Therapeutics, Survey Genomics, NewLimit, Amgen, and Tenaya, owns stock in Arsenal Biosciences, Site Tx, Cellanome, Spotlight Therapeutics, NewLimit, Survey Genomics, Tenaya and Lightcast and has received fees from Site Tx, Arsenal Biosciences, Cellanome, Spotlight Therapeutics, NewLimit, Gilead, Pfizer, 23andMe, PACT Pharma, Juno Therapeutics, Tenaya, Lightcast, Trizell, Vertex, Merck, Amgen, Genentech, GLG, ClearView Healthcare, AlphaSights, Rupert Case Management, Bernstein and ALDA. A.M. is an investor in and informal advisor to Offline Ventures and a client of EPIQ. The Marson laboratory has received research support from the Parker Institute for Cancer Immunotherapy, the Emerson Collective, Arc Institute, Juno Therapeutics, Epinomics, Sanofi, GlaxoSmithKline, Gilead and Anthem and reagents from Genscript and Illumina. P.D.G. is a founder of and has received funding from Juno Therapeutics; is a founder and SAB (scientific advisory board) member of, has equity in, and is receiving research support from Affini-T Therapeutics; is an SAB member of and has equity in Immunoscape, RAPT Therapeutics, Earli, Metagenomi, Nextech, and Catalio; and receives research support from Lonza.

## Abstract

**Background:** Progressive multifocal leukoencephalopathy (PML) is a frequently fatal disease of the central nervous system caused by JC virus (JCV). Survival is dependent on early diagnosis and ability to re-establish anti-viral T cell immunity. Adoptive transfer of polyomavirus-specific T cells has shown promise; however, there are no readily available HLA-matched anti-viral T cells to facilitate rapid treatment.

**Objective:** Identify epitopes of the JCV major capsid protein VP1 that elicit an immune response in the context of human leukocyte antigen allele A*02:01 (HLA-A2) and isolate cognate T cell receptors (TCRs) from healthy donors. Evaluate individual VP1-specific TCRs for their capacity to be expressed in T cells and clear JCV *in vitro*.

**Methods:** PBMCs from HLA-A2+ healthy donors were stimulated with peptide libraries tiled across the JCV VP1 protein. Multiple rounds of stimulation were performed to identify the antigens that induced the largest expansion and CD8^+^ T cell response (measured as INF*γ*, TNF*α*, CD137, and CD69 expression). High-affinity, antigen-specific CD8^+^ T cells were isolated based on intensity of tetramer binding for downstream single-cell TCR sequencing. Candidate TCRs were selected based on tetramer binding affinity and activation assays. Promising TCRs were introduced into the T cell genome via viral transduction for *in vitro* validation including peptide-pulsed K562 cells and astrocyte cells, and JCV-infected astrocytes.

**Results:** Four conserved JCV VP1 epitopes (amino acids 100-108, 251-259, 253-262, and 274-283) presented by HLA-A2 were identified. VP1(100-108) consistently elicited the highest level of IFN-*γ* production from multiple donors and this peptide is in a highly conserved region of VP1. We next identified fourteen high avidity TCRs specific for VP1(100-108). When virally transduced into primary human T cells, seven of these TCRs demonstrated specific binding to VP1(100-108):HLA-A2 tetramers, and four showed increased IFN-*γ* response when incubated with peptide. Primary CD8^+^ T cells expressing two of these TCRs cleared both HLA-A2 positive K562 cells and HLA-A2 positive SVG astrocyte cell line presenting exogenously added VP1 peptide at a range of E:T ratios. In addition, both TCR-transduced T cell populations effectively lysed JCV-infected astrocytes.

**Conclusions:** We identified JCV VP1 epitopes that are immunogenic in the context of HLA-A2 MHC-I, including epitopes that have not been previously described. The VP1(100-108) epitope was used to isolate HLA-A2-restricted TCRs. When cloned into primary human CD8^+^ T cells, these TCRs recognized VP1 (100-108)-presenting targets, and the transduced T cells conferred cytotoxic activity and eliminated K562 and astrocyte cells displaying the VP1(100-108) peptide and not sham peptide, as well as JCV-infected astrocytes. Taken together, these data suggest that JCV VP1-specific TCRs could be appealing therapeutics for HLA-A2+ individuals with PML in whom intrinsic T cell immunity cannot be rescued.

## Introduction

Progressive multifocal leukoencephalopathy (PML) is a devastating brain infection caused by JC virus (JCV) [1]. JCV is a ubiquitous [2] and usually benign DNA polyomavirus that establishes a persistent but asymptomatic infection in many healthy individuals. However, it can reactivate and mutate to become neurovirulent in chronically immunocompromised patients [2]. Patients with HIV/AIDS, idiopathic lymphopenia, sarcoidosis patients who are on chronic immunosuppressive therapies for cancer, solid organ or hematopoietic cell transplant, or autoimmune disease are all at risk of developing PML [3-5]. PML is almost universally fatal if the patient’s immune system cannot be rapidly reconstituted. Even with immune reconstitution, 30-50% of patients die in the first few months, and 70% of survivors are left with ongoing, severe disability [6, 7].

While there are no rigorously proven anti-viral therapies for PML [2], multiple threads of evidence suggest that an active adaptive immune response is sufficient to control the infection. Specifically, restoring T cell immunity quickly is paramount to control the disease. For instance, programmed cell death protein-1 (PD-1) checkpoint inhibitors that boost T cell function have stabilized or led to clinical improvement in select PML patients who had persisting T cell populations at the time of treatment [8-16]. Additionally, JCV-specific CD8+ and CD4+ T cells are always detected in survivors [17]. CD8+ T cells targeting a JCV capsid protein, VP1, play an important role in viral control and recovery [18-21]. CD4+ helper T cell responses are critical for generating and maintaining effective CD8+ T cell responses independent of CD4+ T cell-mediated recognition of infected cells [22].

For PML patients in whom immune reconstitution is not possible, early-stage clinical trials have suggested that adoptive transfer of allogeneic polyomavirus-specific T cells can provide therapeutic benefit [23-29]. However, healthy donor-derived cell-based therapies for adoptive transfer take weeks to months to create. This treatment delay can permit further disease progression, resulting in severe neurologic disability or death. Here, as a first step toward designing an off-the-shelf T cell therapeutic that could be available at the time of PML diagnosis, we comprehensively screened and validated CD8+ T cell JCV VP1 epitopes and their cognate T cell receptors (TCRs) from HLA-A2+ donors, as HLA-A2 is the most prevalent class I HLA in North America [30, 31]. We then assessed the ability of CD8+ T cells expressing these JCV VP1-specific TCRs to clear JCV infection *in vitro*.

## Methods

### Screening of epitopes: Cell stimulation and flow cytometry

Healthy donor frozen leukopaks (STEMCELL) were genetically screened by STEMCELL for HLA-A2 positivity, those with multiple HLA class 1 subtypes were excluded. 1 × 10^8^ peripheral blood mononuclear cells (PBMCs) from six donors were stimulated with peptide pools of a 15-mer peptide library overlapping by 11 amino acids (GenScript) spanning the entire VP1 protein (GenBank: ASV51916.1; sequences deposited on Dryad [32]) which yielded nine peptide pools total (eight pools of ten peptides each and one pool of six peptides). Each donor’s cells were stimulated in Roswell Park Memorial Institute (RPMI) T cell media (Thermo Fisher 11875093), 10% Value HI fetal bovine serum (FBS) (Thermo Fisher A5256801), 0.1% 2-Mercaptoethanol (Life Technologies 21985023), 1% MEM Non Essential Amino Acids Solution 100x (Fisher Scientific 11-140-050), 1% sodium pyruvate 100mM (Life Technologies 11360070),1% penicillin-streptomycin-glutamine 100x (Life Technologies 10378016), and 1% MEM (Vitamin Solution 100x Life Technologies 11120052) supplemented with 1:2000 of 30ug/mL IL-21 (R&D systems 8879 IL 010), 1:2000 of 15ug/mL IL-7 (Biolegend 581902), 1:2000 of 45ug/mL IL-15 (R&D systems 247 ILB 025), and 1:1000 of 200ug/mL IL-2 (Peprotech 200-02). Cells were maintained at 37°C and 5% CO_2_, and half of the media was changed with fresh media containing 2x IL-2 every three to four days.

The cells were re-stimulated every seven to ten days with 1 × 10^6^ irradiated PBMCs (4000 cGy on X-RAD 320) as antigen presenting cells, previously pulsed with corresponding peptides in serum free media for two to three hours. After three rounds of stimulation, the donor T cell response was tested in a T2 co-culture assay [33]. Specifically, HLA-A2+ T2 cells were stimulated with peptides at 10*μ*g/mL for two hours at 37°C and 5% CO_2_ in RPMI. The antigen-loaded T2 cells were subsequently washed to remove excess peptide, and co-cultured with the T cells for six-hours in the presence of protein transport inhibitor 0.2uL BD GolgiPlug (BD Biosciences 555029), 0.2uL GolgiStop(BD Biosciences 554724), and 0.5uL BD FastImmune CD28/CD49d (BD Biosciences 347690) per well. After incubation, cells were first stained with the LIVE/DEAD fixable aqua dead cell stain kit (Invitrogen L34966), followed by anti-CD8 (PE-Cy7, Invitrogen 25-0087-42) stain. Next, cells were fixed and permeabilized using BD Cytofix/Cytoperm kit (BD Biosciences, 554714) and stained with anti-TNF-*α* (FITC, BioLegend 502915) and anti-IFN-*γ* (APC, BioLegend 502516) antibodies. Cells were analyzed using a Fortessa flow cytometer (BD).

Based on the highest stimulatory signals in TNF-*α*/IFN-*γ* assays from multiple donors, selected peptide pools were deconvoluted for individual peptide testing in different donors using the same stimulation protocol to identify the most immunogenic peptides in the pools. In addition to TNF-*α*/IFN-*γ*, cells were stained with antibodies against early activation markers (CD137 (BV410, BioLegend 309820), CD69 (PE, BioLegend 310906)) after 18-24 hours and analyzed with a Fortessa flow cytometer (BD).

Once the most immunogenic individual 15mers were identified, a predictive algorithm (Immune Epitope Database (IEDB) [34]) was used to identify nested 8-10 AA sequences with optimal HLA-A2 binding. These peptides were synthesized (GenScript, purity >95%). To confirm a reproducible TNF*α*/IFN-*γ* response, HLA-A2 PBMCs from six different donors were stimulated with this new 8-10 AA peptide pool in the same manner described above, and T cells from three donors were stimulated to identify the most immunogenic candidate antigen among the group.

Candidate antigen sequences were screened against 749 available JCV genomes recovered from the cerebrospinal fluid or brains of PML patients (NCBI and [35], sequences deposited on Dryad [32]) to ensure that the corresponding VP1 regions were conserved and not mutated away from the wild type JCV VP1 sequence. In addition, candidate antigen sequences were screened in BLASTx to assess for similarity to human proteins as a means of assessing for the potential of cross-reactive responses resulting in autoreactivity.

### T cell line sorting and TCR Sequencing

Candidate antigen-stimulated T cell lines from three donors were pooled and stained with a viability dye (Thermo Fisher 65-0863) and split into three fractions for subsequent staining and sorting. Each fraction was stained with a different concentration (1:400, 1:2000, 1:4000) from 0.4mg/ml stock of JCV VP1(100-108):HLA-A*02:01 tetramer (Fred Hutchinson Cancer Center Tetramer Core) for 30 min at 4°C, followed by staining with anti-CD3 and anti-CD8 (BD Biosciences). The use of limiting dilutions of the tetramer allowed for normalization for the level of TCR expression, to help avoid isolating T cells that exhibit high intensity tetramer binding based primarily on high levels of TCR expression rather than expression of a high affinity TCR [36]. On the sorter (Aria II, BD Biosciences), live CD3+ CD8+ T cells were further gated into “tetramer negative”, “tetramer positive”, and “top tetramer positive” (corresponding to ∼10% of the tetramer+ population) to identify negative clones, VP1-specific clones, and VP1-specific clones with higher TCR affinity, respectively.

Cells from each tetramer gate were sorted from each sample fraction, DNA was extracted, and bulk TCR beta libraries were prepped and sequenced on an Illumina MiSeq instrument (Adaptive ImmunoSeq, hsTCRb kit) [37] to quantify relative TCR frequency in each gate/sample fraction. TCR clones were defined based on V, D, J gene usage and TCRb CDR3 amino acid sequences using ImmunoSeq Analyzer 3.0 (Adaptive Biotechnologies).

After sorting sufficient cell numbers for bulk TCRb sequencing, 1:400 and 1:4000 tetramer dilution positive group were sorted again for single cell sequencing (10X Genomics, Chromium Single Cell V(D)J Enrichment Kit, Human T Cell, PN-1000005). cDNA libraries were prepped and sequenced on an Illumina NovaSeq 6000 instrument with a goal of 50,000 paired-end reads/cell with read lengths of 26 base pairs (read 1) and 90 base pairs (read 2). Cellranger 6.1.2 and Loupe VDJ Browser 4.0 (10X Genomics) were used to identify paired TCRa and TCRb chains and extract V, D, J and CDR3 amino acid sequences for subsequent TCR assembly.

### Identifying candidate TCRs

TCR clones that were statistically significantly enriched in the “top tetramer positive” gate relative to the “tetramer negative” and “tetramer positive” populations and were abundant in the scTCR dataset were presumed to represent most of the high affinity TCRs. These were selected for further arrayed validation as true high affinity TCRs. These top candidates were cloned into the pRRLSIN lentiviral vector as previously described [38] and used to transduce primary human CD8+ T cells. Specifically, CD8+ T cells were isolated by negative selection from two healthy donor PBMCs (EasySep kit, STEMCELL), activated for 4-24h with anti-CD3/CD28 Dynabeads (Thermo Fisher 11161D) in the presence of 50IU/ml IL-2, and spinfected (1000 x g, 90 min, 30°C) with lentiviral supernatant and polybrene (5*μ*g/ml). Cells were maintained in culture for up to 10 days prior to functional assessment, with fresh media and cytokine addition every 2 days (IL-2 at 50IU/ml, IL-7 at 2250 IU/ml, IL-15 at 2250 IU/ml). These cells were then tested for IFN-*γ* production and tetramer binding (at 1:200) as described above. TCRs with both persistent function and binding were selected for further validation.

### Cell Killing assay

Primary human CD8+ T cells from 2 donors were pooled and stimulated with anti-CD3/anti-CD28 magnetic beads and transduced with lentiviral vectors encoding the indicated TCRs. After 5 days of bead stimulation, T cells were separated and rested 7-10 days prior to assay. TCR expression was validated by tetramer staining. 30,000 K562 cells (ATCC CCL-243) expressing HLA-A2 and BFP were added to each well of a round bottom 96 well plate with 150uL T cell media (as described above). Each peptide was added at 1uM at time 0. T cells were added at the indicated E:T ratio 2 hours later after a wash. In addition, a TCR (DMF5) targeting MART1 antigen (ELAGIGILTV) was used as a specificity control (“sham”) [39]. K562 viability was quantified by flow cytometry by gating on BFP-positive viable cells per input volume after 72 hours of co-culture. Viable cell count was based on the number of viable K562 cells per input volume normalized to the average of the condition without T cells for each peptide.

### Infection of SVG Astrocyte Cells

HLA-A2 SVG astrocyte cells (generously provided by Walter Atwood, Brown University) were plated at 50-60% confluency in T-75 flasks (Corning 430641U) in T cell media 24 hours prior to infection. The media was removed, and cells were washed with warmed PBS at 37°C. The cells were infected with MAD-4 JCV (ATCC VR-1583, identical VP1 sequence as our source peptide), at a multiplicity of infection (MOI) of 0.2 with 1580uL of infectious media (DMEM with 2% FBS) in drop wise fashion throughout the flask or with 50ug of VP1(100-108) peptide. Cells were incubated for two hours at 37°C in a humidified incubator with 5% CO_2_, with intermittent shaking every 15 minutes. Afterwards, the media was replaced with T cell media.

*Mycoplasma pneumoniae* testing was performed on the media following the ATCC 166553 protocol.

### Assessment of T cell populations to clear JCV infection in vitro

JCV-reactive TCR-transduced human CD8+ T cells or sham T cells were added to infected SVG cells at an E:T ratio of (5:1-1:10), and cells were observed for 72 hours in co-culture. T cells were isolated and evaluated by flow cytometry for CD137 activation markers. SVG astrocytes were also monitored for viability (GFAP, FITC ThermoFisher Scientific 53-9892-82).

### Digital droplet PCR of SVG cells and media supernatant

DNA was extracted from the supernatant and the astrocyte cells after co-culture with T cells at 72 hours (QIAGEN QIAamp MinElute Virus Spin Kit) according to manufacturer’s instructions. DNA was eluted in 30uL of nuclease-free water, and the DNA concentration was measured using a NanoDrop ND-1000 spectrophotometer. Based on a previously published protocol [40], DNA was digested in a digest mix at a 1:1 ratio (15uL sample, 15uL digest mix). The digest mix contained the following: 3.65uL nuclease-free water, 1uL 10x NEB Buffer, 0.1uL 2% BSA, and 0.25uL HindIII restriction enzyme. Samples were incubated on a shaker at 37°C for 30min at 300rpm. Primers and probe were created using the VP1 sequence of JC Virus on Primer3Plus and ordered from IDT with primers at a 25nM concentration and probe at 250nM concentration with a 5’ FAM. Forward primer: GAGGCAGCAAGCAATGAATC, Reverse Primer: GCCAAACAAAGGGTTGACAG, Probe: AGGCCACCCCAGCCATATAT. DdPCR master mix was made in duplicate containing 12.5uL ddPCR supermix for probes, 1.25uL 20x primer-probe mix (100uL of 20x primer-probe mix created by adding 18uL 100uM forward primer, 18uL 100uM reverse primer, 5uL 100uM probe, and nuclease-free water added to bring total volume up to 100uL). Following this, 30uL of master mix was aliquoted into a PCR tube, and 20uL of 1:2.5 diluted digested DNA was added. All reagents were vortexed well before droplet generation.

In a DG8 cartridge, 20uL of the DNA ddPCR mixture was added into each sample well, and 70uL of droplet generation oil for probes was added into the oil well. Once droplets were generated, they were dispensed into ddPCR 96 well semi-skirted plates. The plate was heat sealed with foil and placed into the thermocycler. The thermocycler was set with a 2°C/s ramp rate with the following cycling conditions: 95°C 10min for 1 cycle, 94°C 30sec for 40 cycles, 56°C 60sec for 40 cycles, 98°C 10 min for 1 cycle, 4°C hold. Plate was then placed into a QX 200 Droplet Reader for probe detection.

### Patient sample flow cytometry

The patient sample was thawed and stained with live dead aqua (Invitrogen L34966), anti-CD8 (PE-Cy7, Invitrogen 25-0087-42), and JCV VP1(100-108):HLA-A*02:01 (PE) (and the bioequivalent BKV VP1(100-108): HLA-A*02:01 (APC) tetramer as described above, followed by analysis on a Fortessa flow cytometer using FACSDiva software (BD). Data were analyzed using FlowJo software (BD).

### Statistical Analysis

For all statistical analyses, one-way ANOVA testing, t-tests, and EC50 calculations were completed on GraphPad Prism 9 software.

## Results

### Identification of immunogenic epitopes from the VP1 protein

PBMCs from six HLA-A2 donors were individually stimulated with nine overlapping peptide pools spanning JCV VP1. After three rounds of stimulation every 7-10 days, the cells were stained for evidence of cytokine production (IFN-*γ* and TNF-*α*) (Figure 1A). Peptide pools 3 and 7 produced the most robust stimulation in terms of the frequency of responding T cells (Figure 1B). PBMCs from four HLA-A2 donors were then stimulated with 10 individual peptides from pools 3 and 7 (Supplemental figure 1) to again look for cytokine production and early activation markers (CD137 and CD69).

**Figure 1:**
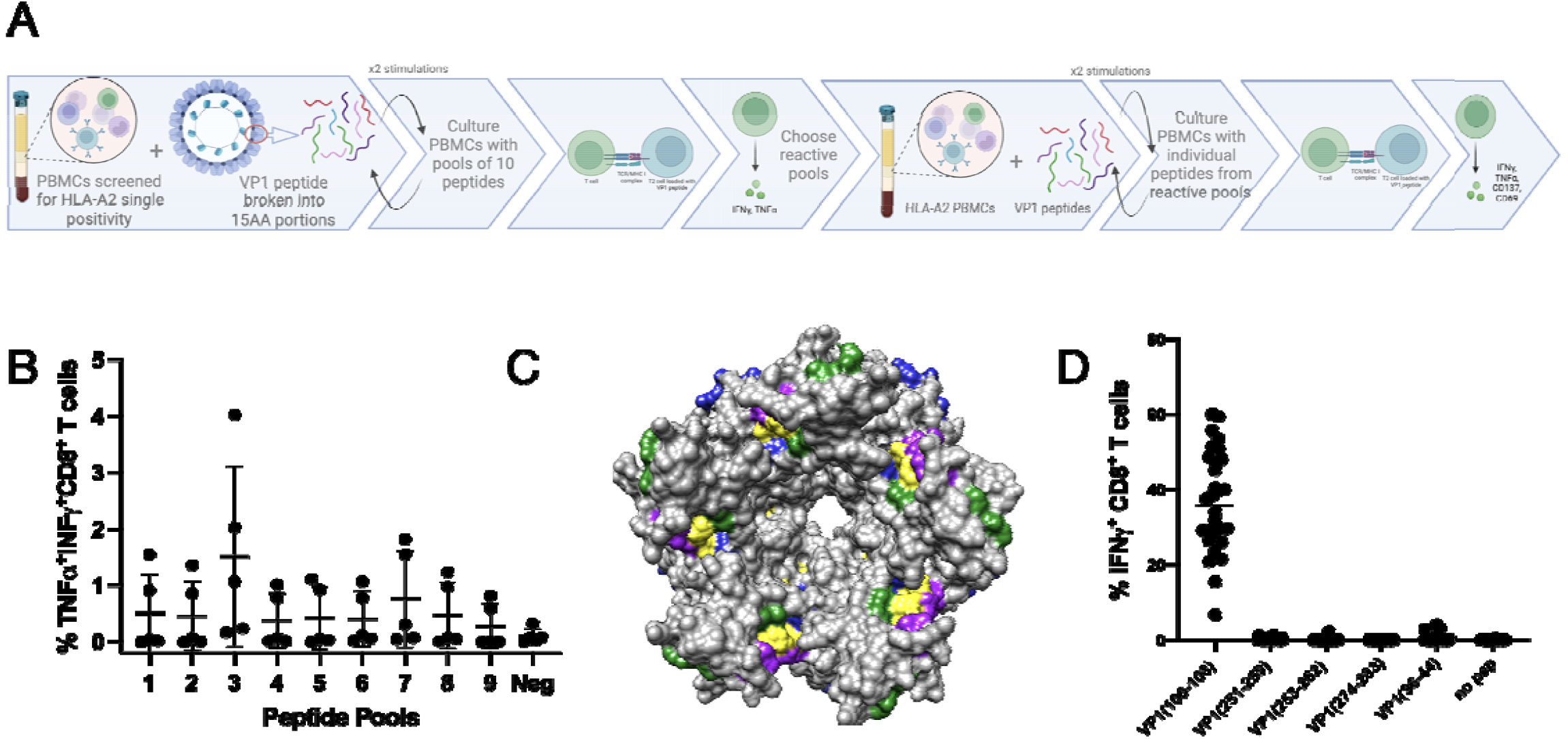
Defining immunogenic peptide. **A**. Schematic of experimental layout. **B**. CD8+ T cells from 5 donors stimulated 3x with pools of 10 peptides, except pool 9 which had 6 peptides. **C**. 3D modeling of VP1 pentamer highlighting epitopes (Green: VP1 (100-108), Yellow: VP1 (251-259), (253-262), (274-283), Purple: Published epitopes, Blue: Predicted epitopes from IEDB. **D**. CD8+ T cells from 3 donors, 10 cell lines each, stimulated 3x with pools of peptides and then individual T2 IFN-*γ* response.

The peptides that induced the greatest cytokine production and expression of activation markers across the A2 donors clustered into 3 different regions of the VP1 protein (pool 3: peptides 24-26; pool 7: peptides 62-64 and 68-69) (supplemental figure 1). Using IEDB predictive MHC binding algorithms, four 8-10mer peptides from these VP1 regions were identified, 2 of which overlapped: VP1: amino acids 100-108, 251-259, 253-262, and 274-283. Three of these epitopes were distinct from previously described epitopes [20, 21, 41]. Comparative analysis of previously published epitopes [20, 21, 41], predicted epitopes (IEDB), and the epitopes we identified, revealed VP1 (100-108) to be found among all 3 groups. Epitopes from all these categories clustered in one region of the pentameric VP1 (Figure 1C).

The four predicted VP1 epitopes (100-108, 251-259, 253-262, and 274-283) within the three VP1 regions we identified as well as three previously published epitopes, (VP1(36-44) [20], VP1(117-123) and VP1(288-294) [41], were selected for further testing. The 7 peptides were tested in six donors individually (except for VP1(117-123) and VP1(288-294) which were tested in two donors) for inducing IFN-*γ* and TNF-*α* production and a statistically significant difference was detected between 5 of 7 of the peptides and no peptide (unpaired t test, p <0.05) but not between each of the five peptides (Supplemental figure 2). Two of the peptides VP1(251-259) and VP1(228-294) did not elicit a statistically significant increase in IFN-*γ* and TNF-*α* production compared to no peptide. To determine the most immunogenic peptide, five of the peptides were pooled to stimulate PBMCs from three new HLA-A2 donors. After three rounds of stimulation, the cells were then tested with each individual peptide from the pool of 5. Based on the frequency of responding T cells, VP1(100-108) was the most immunogenic (Figure 1D).

The VP1(100-108) sequence was next assessed in 749 published JCV genomes from the brains and cerebrospinal fluid of PML patients. The VP1(100-108) amino acid sequence was conserved in >99.9% of those sequences (supplemental figure 3). Furthermore, the VP1(100-108) amino acid sequence was not homologous with any human protein based on a BLASTx search.

### Isolation of JCV VP1(100-108)-reactive TCRs

Three sets of donor PBMCs were stimulated with VP1(100-108) for three rounds. Stimulated T cells were pooled and stained with VP1(100-108) HLA-A2 tetramers at limiting dilutions and then sorted for bulk TCR sequencing and scTCR sequencing. 36,941 TCR beta sequences were identified from the bulk TCR sequencing dataset and ranked by fold enrichment in the “top tetramer positive” sorting gate in comparison to the total “tetramer positive” and “tetramer negative” populations (Supplemental figure 4A). 246 TCRs demonstrated >3-fold enrichment in the “top tetramer positive” gate.

Approximately 130,000 T cells were submitted for scTCR sequencing at both 1:400 and 1:4000 tetramer dilutions, identifying 2,048 and 2,148 unique TCR pairs, respectively. 126 of these TCR pairs matched clones demonstrated >3-fold enrichment in at least one of the stringent tetramer+ gates by bulk TCR sequencing. From this set, we selected the fourteen TCRs displaying highest tetramer avidity based on the bulk TCR beta datasets (Figure 2B, supplemental figure 4B). These TCRs were then lentivirally transduced into primary human CD8+ T cells from two different donors to observe both tetramer binding and functional IFN-*γ* assays (Figure 2C). TCR3 and TCR8 demonstrated robust functional response across both donors with varied avidity were selected for further evaluation.

**Figure 2:**
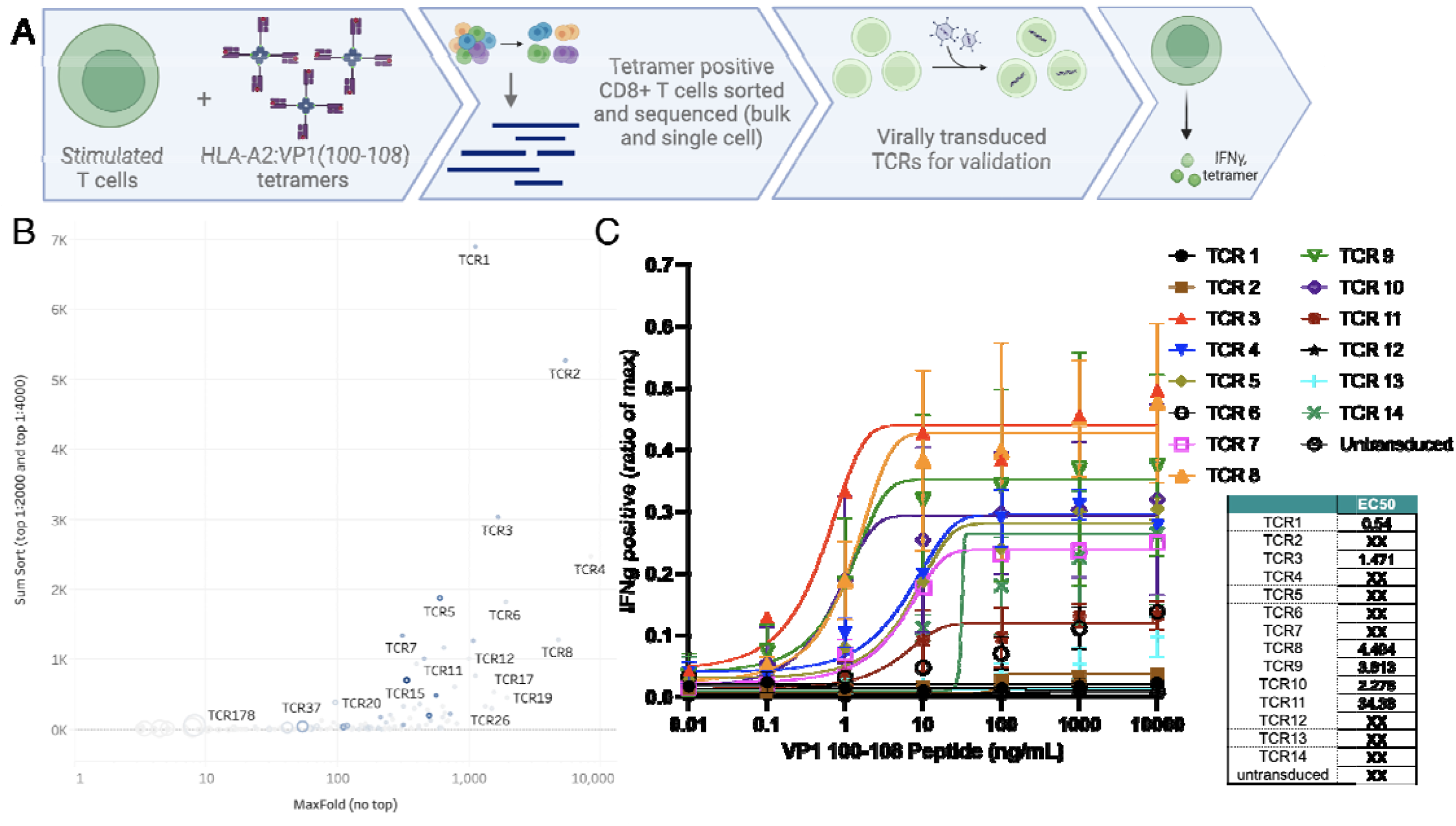
JCV VP1 (100-108) specific T cell receptors (TCR). **A**. Schematic of experimental layout. **B**. Pooled CD8+ T cell lines from 3 healthy donors were split into three different fractions and stained with low, intermediate, or high tetramer concentration. From samples stained with low and intermediate tetramer concentration, live CD3+ CD8+ T cells were sorted from the 90th percentile tetramer(+) gates. Shown here is the cumulative clone count in the 90th percentile gates (y-axis) plotted against maximum enrichment score out of the 11 calculated for each clone (x-axis). Symbol size corresponds to clone count in tetramer-negative gate. Symbol color indicates clone abundance in the single cell TCR dataset (low tetramer concentration condition). High affinity TCRs have high cumulative clone count, high maximum enrichment score, and small symbol size. SumSort = total number in top 10% tetramer binder gate (i.e. 90th percentile) in the 1:2000 + 1:4000 tetramer staining condition. MaxFold = greatest value for fold enrichment (out of the calculated 11). **C**. Top 14 T cell receptors were virally transduced into human primary T cells in 2 donors. TCR-transduced cells were stimulated with different amounts of VP1(100-108) peptide and tested for intracellular INF-*γ* content as ratio of positive control (max). EC50 (M) scores populated in tabular format.

### Functional validation of VP1(100-108) specific TCRs

CD8+ T cells were first transduced with either TCR 3 or 8 (Supplemental figure 5) and co-cultured with HLA-A2 K562 cells pulsed with VP1(100-108) or a non-specific peptide. After 48 hours, the transduced T cells mediated ∼40-50% killing of the targets at an E:T of 0.5, and 100% killing at an E:T of 2:1, but not the non-specific peptide (Figure 3B). TCR3 and TCR8 T cells were then co-cultured with SVG astrocyte cell lines pulsed with VP1(100-108) peptide at varied E:T ratios. They upregulated expression of the CD137 activation marker and cleared up to 60% of astrocytes in comparison to no T cell (<1% clearance) and sham T cell (∼15% clearance) controls (Supplemental figure 6).

**Figure 3:**
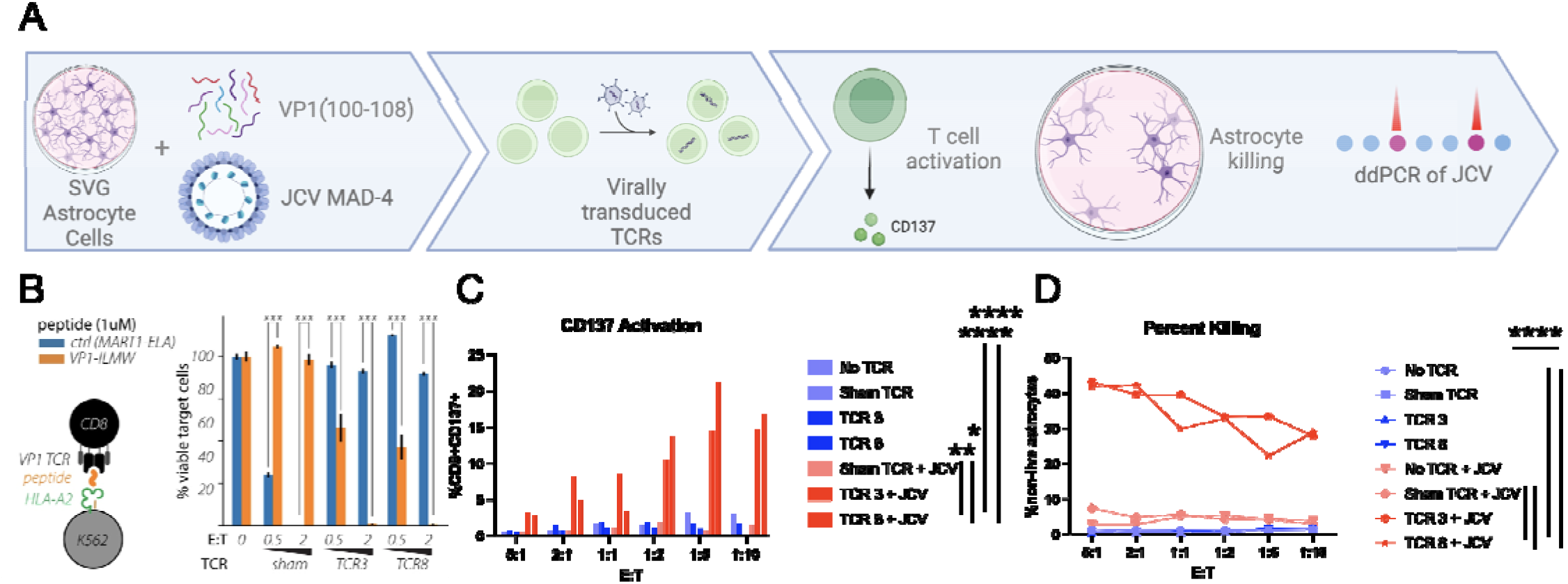
Validation of JCV VP1 (100-108) T Cell Receptors (TCRs). **A**. Schematic of experimental layout. **B**. TCR 3 and TCR 8 tested against K562 presenting VP1(100-108) on HLA-A2 vs a control TCR (DMF5) with its corresponding peptide (MART1) after 48h incubation at unsorted E:T of 0.5 and 2. Error bars repres nt SEM for 3 technical replicates from one representative experiment. p<0.001 by unpaired T test. **C**. After 72 hours of co-culture with JCV-infected SVG cells and no T cells (no TCR), sham TCR, TCR 3, and TCR 8 looking at live CD8+ T cell CD137 positivity via flow cytometry. TCR 3 and 8 vs no TCR p value <0.001; TCR 3 vs sham TCR+JCV p=0.0032 and TCR 8 vs sham TCR+JCV p=0.029 **D**. Percent killing of SVG cells determined by flow cytometry based on gating of loss of live astrocytes. p value <0.001; TCR 3 vs sham TCR+JCV p<0.001 and TCR 8 vs sham TCR+JCV p=0.002 by unpaired T test. Done in duplicate at various unsorted E:Ts, mean results shown.

CD8+ T cells expressing either TCR 3 or 8 were next co-cultured with SVG astrocytes directly infected with JCV, again demonstrating significant CD137 expression activation and efficient clearance of JCV-infected astrocytes (average 34%, between 28-43%) at all E:T ratios compared to no TCR (p value <0.001 for TCR 3 and 8) or sham T cell controls (p value 0.0032 (activation), <0.001 (killing) for TCR 3; p=0.029 (activation), 0.002 (killing) for TCR 8 (Figure 3C-D). This is compared to the direct cytotoxicity of the virus which resulted in death of SVG astrocytes from 50-75% compared to uninfected controls after a mean of 12 days (ranging 10-15 days) (data not shown). In addition, there was a statistically significant reduction in JCV copy number in cell co-cultured with JCV reactive TCR transduced T cells with an average of 2210 copies/uL (633.5-3468 copies/uL) compared to sham T cells with an average of 3263.5 copies/uL (1925-3959 copies/uL); p value 0.035 for TCR 3 vs sham and p=0.0061 for TCR 8 vs sham (supplemental figure 7)).

### Endogenous response to VP1(100-108) peptide in a PML survivor

Using JCV VP1(100-108) tetramers and the equivalent BK virus epitope, we stained PBMCs from an HLA-A2 PML survivor in comparison to an HLA-A2 JCV antibody negative donor. The PML survivor demonstrated robust binding to both tetramers consistent with the presence of a VP1(100-108) reactive CD8+ T cell population that was not identified in the JCV antibody negative donor (Figure 4).

**Figure 4:**
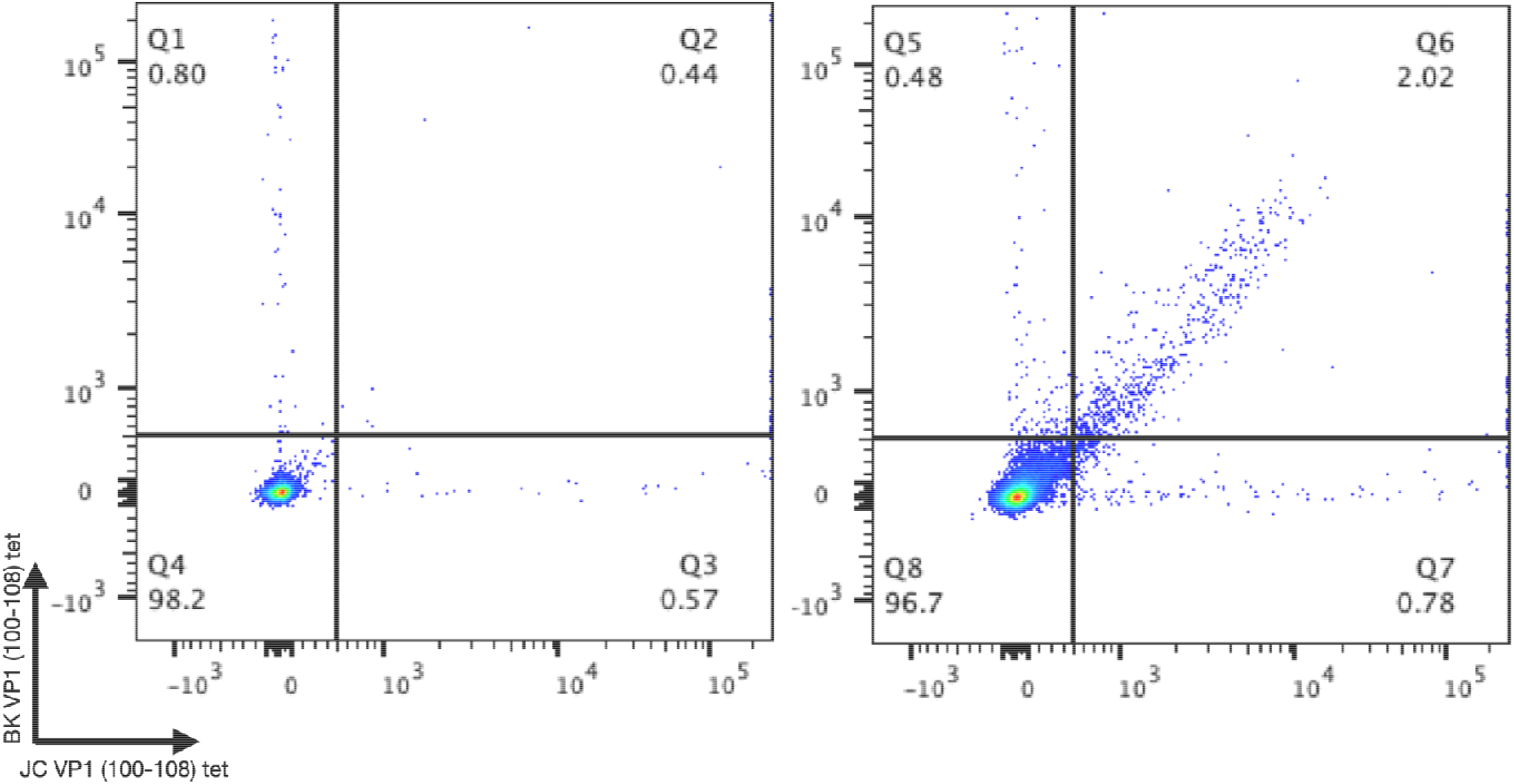
Patient response to JCV and BKV VP1 (100-108). Flow cytometry data of live CD8+ T cells stained with HLA-A2:BK VP1 (100-108) and HLA-A2:JV VP1 (100-108) at 1:50 tetramer dilution with an HLA-A2 positive JCV antibody negative healthy donor on the left and an HLA-A2 confirmed PML survivor on the right.

## Discussion

PML is a devastating neurologic infection that is often fatal without rapid immune reconstitution. Urgent immune restoration to counter the swift spread of JCV throughout the brain is not currently possible for many patients. Here, we identified novel JCV VP1 antigens for individuals with an HLA-A2 background, establishing VP1(100-108) as the most immunogenic epitope. VP1(100-108) is a highly conserved region of the JCV VP1 protein, making it unlikely that JCV will escape from peptide-specific T cells by mutagenesis. We did not find that this amino acid sequence had similarity to human protein sequences, and it is recognized by TCRs from multiple healthy donors we evaluated, suggesting that T cells specific for this antigen in this dominant HLA-A2 background would have low potential for inducing autoimmunity. Combining bulk TCR sequencing and single cell analysis, we further identified paired TCR sequences reactive for this therapeutically attractive antigen. We demonstrate that primary human CD8+ T cells engineered with these TCRs efficiently clear VP1(100-108)-presenting cells and cells infected with wild type JCV. Finally, we found T cells reactive against this antigen in a PML survivor.

The JCV VP1 protein covers most of the external surface of the JCV capsid, creating a depression required for receptor binding of sialylated glycans and subsequent entry of the virus into a host cell [42]. Thus, this protein is vital for ongoing JCV infectivity. In addition, VP1 is a highly conserved protein, with only rare point mutations that change the virulence of the virus [35, 43, 44]. VP1 has been reported to be a prominent target of effective host cellular immunity [18-21]. Indeed, PML survivors have an abundance of VP1-specific T cells, and response to VP1 peptides has been associated with better outcomes in PML patients [21].

There have been only three studies exploring the antigen specificity of the CD8+ T cell response to VP1 in the past 20 years. Koralnik and colleagues also identified VP1(100-108) as an immunogenic target after screening 24 HLA-A2 patients with PML, and evaluating potential responses to 11 predicted epitopes across multiple JCV proteins (T, t, VP1, VP2, VP3, and agnoprotein) [21]. They found tetramer binding of CD8+ T cells to VP1(100-108) in 5 out of 7 survivors compared to zero of six non-survivors. Du Pasquier and colleagues assessed the immunogenicity of the whole VP1 protein amongst 36 HLA-A2 PML patients (24 with proven PML) looking for tetramer binding and lysis. They found VP1(36-44) tetramer positive CD8+ T cells in 10 of 11 survivors and only in 1 of 11 patients with progressive PML [20]. A more recent study by Mani and colleagues found two VP1 epitopes: 117-123 and 288-294 [41]. The additional epitopes we identified cluster spatially with the previously identified epitopes on the VP1 pentamer. Our work highlights the importance of defining epitopes by both MHC binding (predicted epitopes), but also by T cell functional studies by looking at both cytokine production (TNF-*α*/IFN-*γ*) and activation (CD137/CD69) of the cells. For example, we found that the VP1(117-123) and VP1(288-294) epitopes defined by Mani and colleagues elicited some CD137 positivity but not a functional response, suggesting that these are less attractive therapeutic targets.

In the 1990s, it was noted that post-transplant patients with Epstein-Barr virus (EBV)-associated lymphoproliferative disorders had better outcomes if they were also given donor T cells that could enhance EBV immunity [45]. This led to treatment of post-transplant or immunosuppressed patients with allogenic, virus-specific T cells for EBV [46], cytomegalovirus, human herpesvirus 6, and even BK polyomavirus [47]. As a result, “off-the-shelf” banks of the virus-specific T cells now exist for individuals with various HLA backgrounds. Moreover, the efficacy of these treatments has sparked interest in utilizing virus-specific CTLs for tackling virus-associated cancers such as those linked to EBV [48, 49], hepatitis B virus [50], and others.

Similarly, the long-term goal of this work is to identify virus-specific TCRs that might serve as the backbone for a PML cellular therapeutic, allowing for the field to move beyond less specific adoptive transfer trials with stimulated polyclonal T cells [23-29]. However, these donor cells require some degree of HLA matching to prevent rejection despite transfer into PML patients who are immunodeficient, with the majority having represented 5/10 MHC alleles or more (though has been reported with as low as 2/10 with success). Rapid advances in clinic-grade targeted gene editing in both autologous and allogeneic T cell therapy settings[51, 52] provide a clearly charted path for efforts (beyond the scope of this paper) to promote graft survival and mitigate graft-versus-host disease, and augment its potency. [53]. These modifications will include disrupting expression of the endogenous TCR chains and modifications to prevent T cell rejection and promote retention (i.e., downregulating beta-2-microglobulin and CIITA; upregulating HLA E and CD47). With such changes, if the host response does become able to reconstitute enough immunity to reject our product, likely there would also be a robust enough recovered adaptive response to control the JCV. Furthermore, a potential superiority of this type of product design is the improved predictability of the outcome as opposed to *ex vivo* stimulated cells that can have mixed and unpredictable outcomes.

This work expands and validates available HLA-A2 restricted epitopes of the JCV VP1 protein, a viral protein critical to the neuropathogenesis of PML. The TCRs that target these epitopes are promising components of a future anti-viral T cell therapeutic targeting viral components in class I HLA-A2 that we are developing and could be rapidly deployed in PML patients for whom timely immune reconstitution is not possible or is insufficient, or in patients for whom more general immune reconstitution might be deleterious (i.e., solid organ transplant patients). The final product would potentially include multiple epitopes across multiple viral proteins with a polyclonal T cell population having varied affinities We further hope to expand this beyond HLA-A2, to other regional dominant HLAs including A1, A3, A11, and A30 to support the diverse patient population afflicted with this disease and not limit it to those who are able to find a closely matched donor.

## Acknowledgements

We would like to thank the participants of the PML natural history trial (NCT01730131) from which we received the patient sample. We would like to thank Dr. Walter Atwood (Brown University) for meaningful regents and fruitful discussions. We would like to thank Dr. Joseph Sabatino (UCSF) for guidance in assay development and ongoing discussions. We would like to thank Collin Spencer (UCSF) for guidance in equipment optimization and interpretation of certain data. In addition, we would like to thank Dr. Stephen Hauser (UCSF) for this unwavering support and Dr. Avindra Nath (NINDS) for his enthusiasm in unifying the initial group. Schematic figures (figure 1A, 2A, and 3A) were created in BioRender. Gupta, S.

