## Supplemental Figures for "Antigen-Specific T Cell Receptor Discovery for Treating Progressive Multifocal Leukoencephalopathy"

**Supplemental Figure 1:** Live CD8+ T cells from 4 donors stimulated 3x from individual peptides from pool 3 and 7 looking at CD137+ and CD69+ or INF-γ+ and TNF-α+ double positive. Error bars represent SEM.

**Supplemental Figure 2:** CD8+ T cells from 6 donors stimulated 3x from individual epitopes looking at INF-γ+ and TNF-α+ double positive responses. p values based on unpaired T test.

**Supplemental Figure 3:** 749 published PML sequences covering the 354 amino acids of JCV VP1 overlapped on each other, with black showing conservation and epitopes discussed bracketed in red.

**Supplemental Figure 4:** A. Schematic flow gating for adaptive sequencing. Pooled CD8+ T cell lines from 3 healthy donors were split into three different fractions and stained with low, intermediate, or high tetramer concentration. **B.** Live CD3+ CD8+ T cells were sorted from tetramer(+) gate from the sample stained with intermediate tetramer concentration. Shown here is clone fold enrichment in tetramer(+) gate (intermediate concentration) over tetramer(-) gate (high concentration) (y-axis) versus clone abundance in the tetramer(+) 90th percentile gate (intermediate concentration). C. Live CD3+ CD8+ T cells were sorted from tetramer(+) gate from the sample stained with low tetramer concentration. Shown here is clone fold enrichment in tetramer(+) gate (low concentration) over tetramer(-) gate (high concentration) (y-axis) versus clone abundance in the tetramer(+) 90th percentile gate (low concentration). Symbol size indicates the statistical likelihood (p-value) of clone enrichment in the most stringent tetramer(+) gate compared to least stringent tetramer(+) gate. Symbol color indicates clone abundance in the single cell TCR dataset (low tetramer concentration condition). High affinity TCRs have high fold enrichment scores and high abundance and are represented with smaller symbol sizes.

**Supplemental Figure 5:** Virally transduced CD8+ primary human T cells were stained with HLA-A2:BK VP1 (100-108) and HLA-A2:JCV VP1 (100-108) at 1:200 tetramer dilution to measure transduction efficiency.

**Supplemental Figure 6:** Virally transduced CD8+ primary human T cells cultured alone or with SVG cells incubated with VP1(100-108). A. CD137 positivity assay done in duplicates at various unsorted E:Ts. No TCR vs TCR 3 and 8 p <0.0001; sham T cells vs TCR 3 p value 0.0039, vs TCR 8 0.0013 B. Percent killing of SVG cells done in duplicates at various unsorted E:Ts.

**Supplemental Figure 7:** ddPCR of JCV infected SVG cells (SVG) or their supernatant (Sup) comparing Sham T cells to VP1 directed T cells (TCR 3 and TCR 8) after 72hour of co-culture. p value 0.035 for TCR 3, SVG vs Sham, SVG and p=0.0061 for TCR 8, SVG vs Sham, SVG based on unpaired T test. ddPCR=droplet digital polymerase chain reaction; TCR=T cell receptor

**Supplemental Figure 1:**

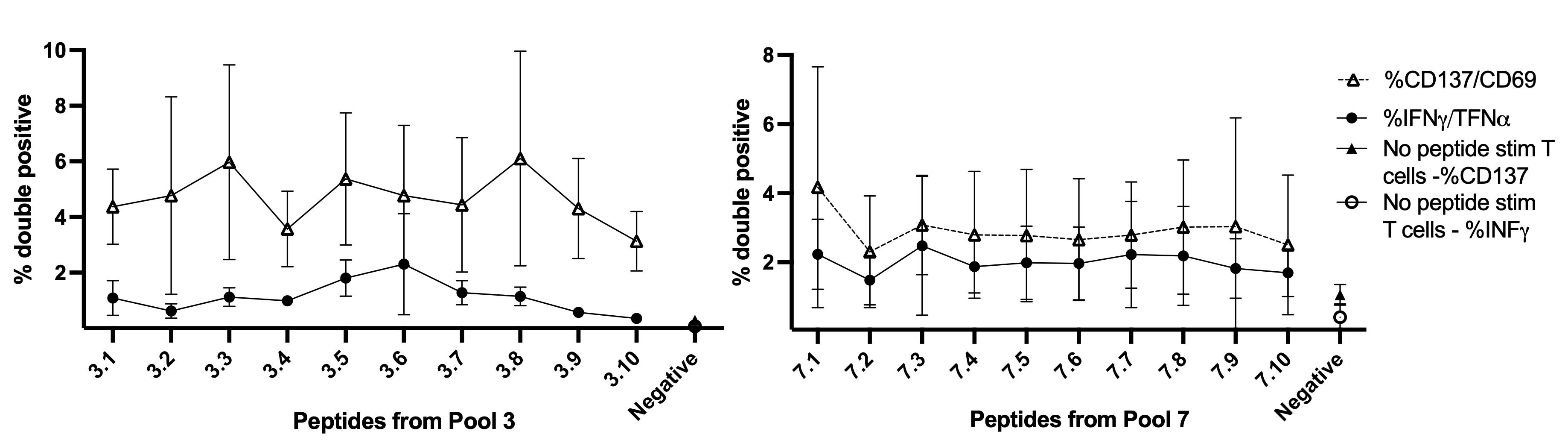

**Supplemental Figure 2:**

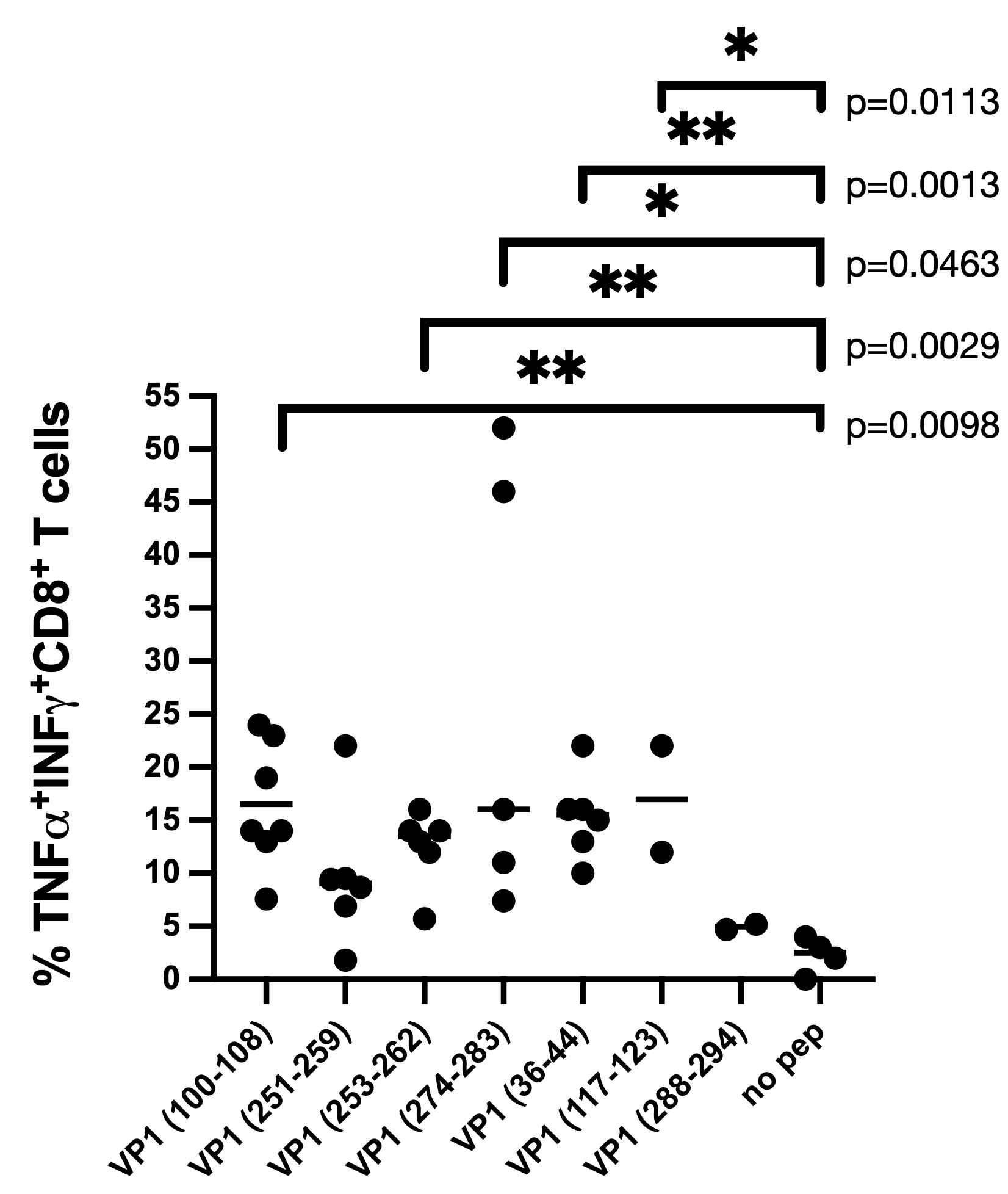

**Supplemental Figure 3:**

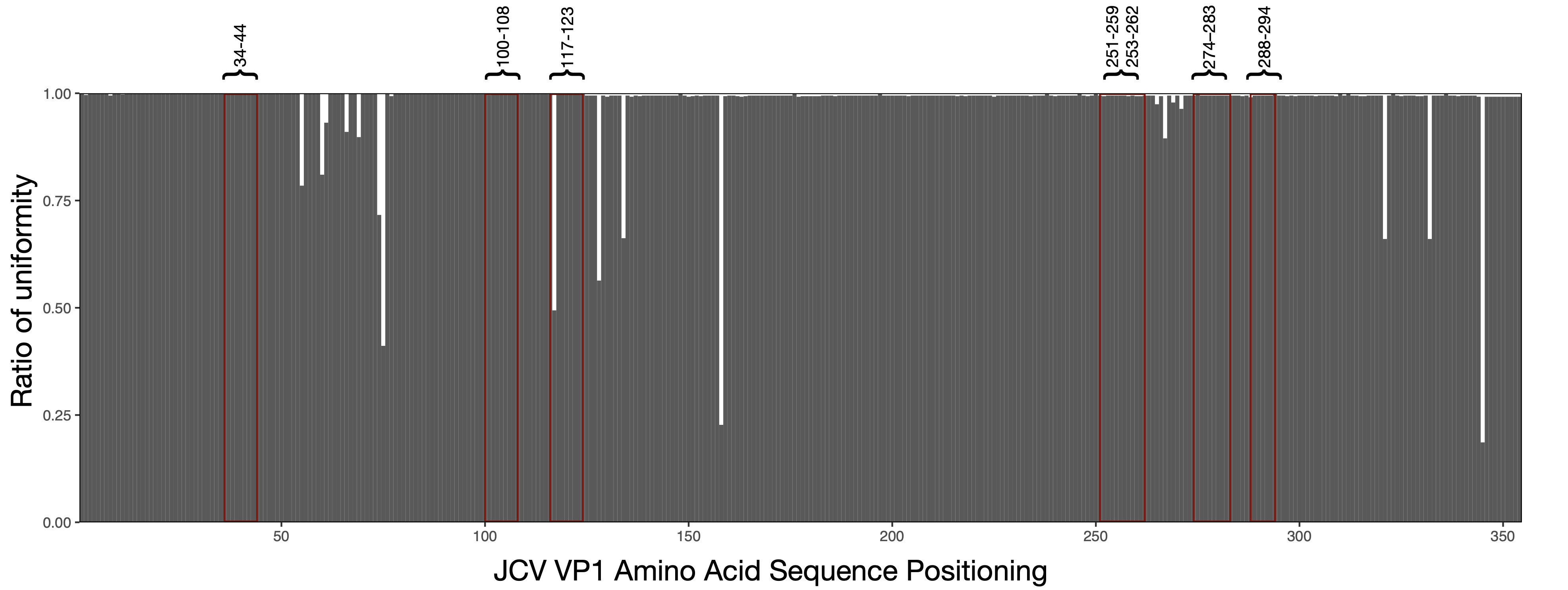

**Supplemental Figure 4:**

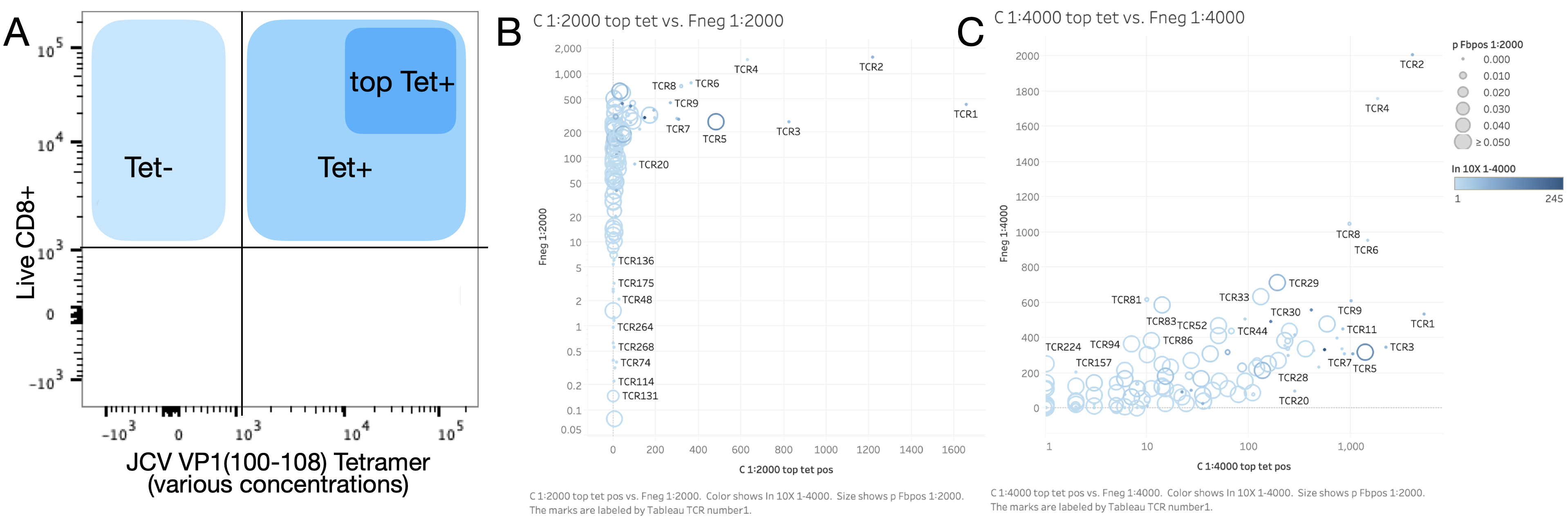

**Supplemental Figure 5:**

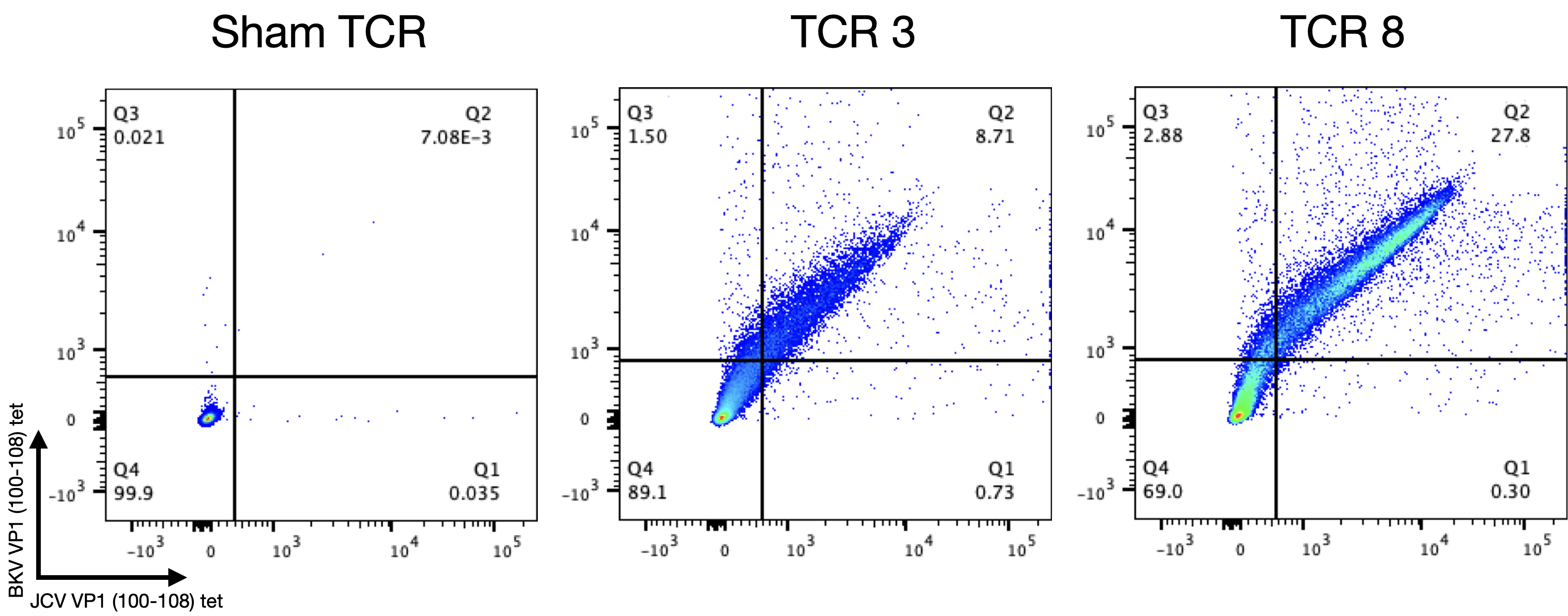

**Supplemental Figure 6:**

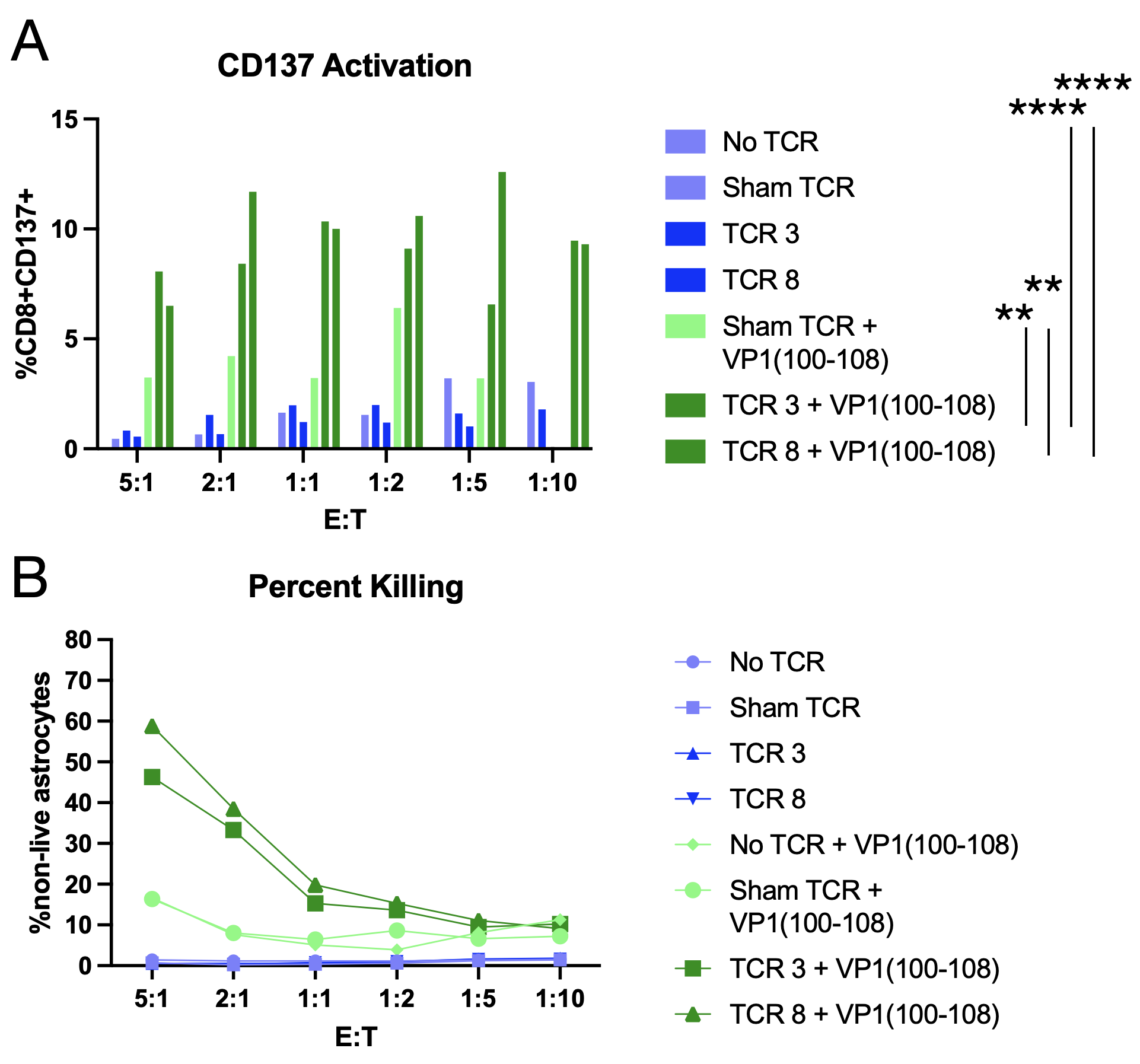

**Supplemental Figure 7:**

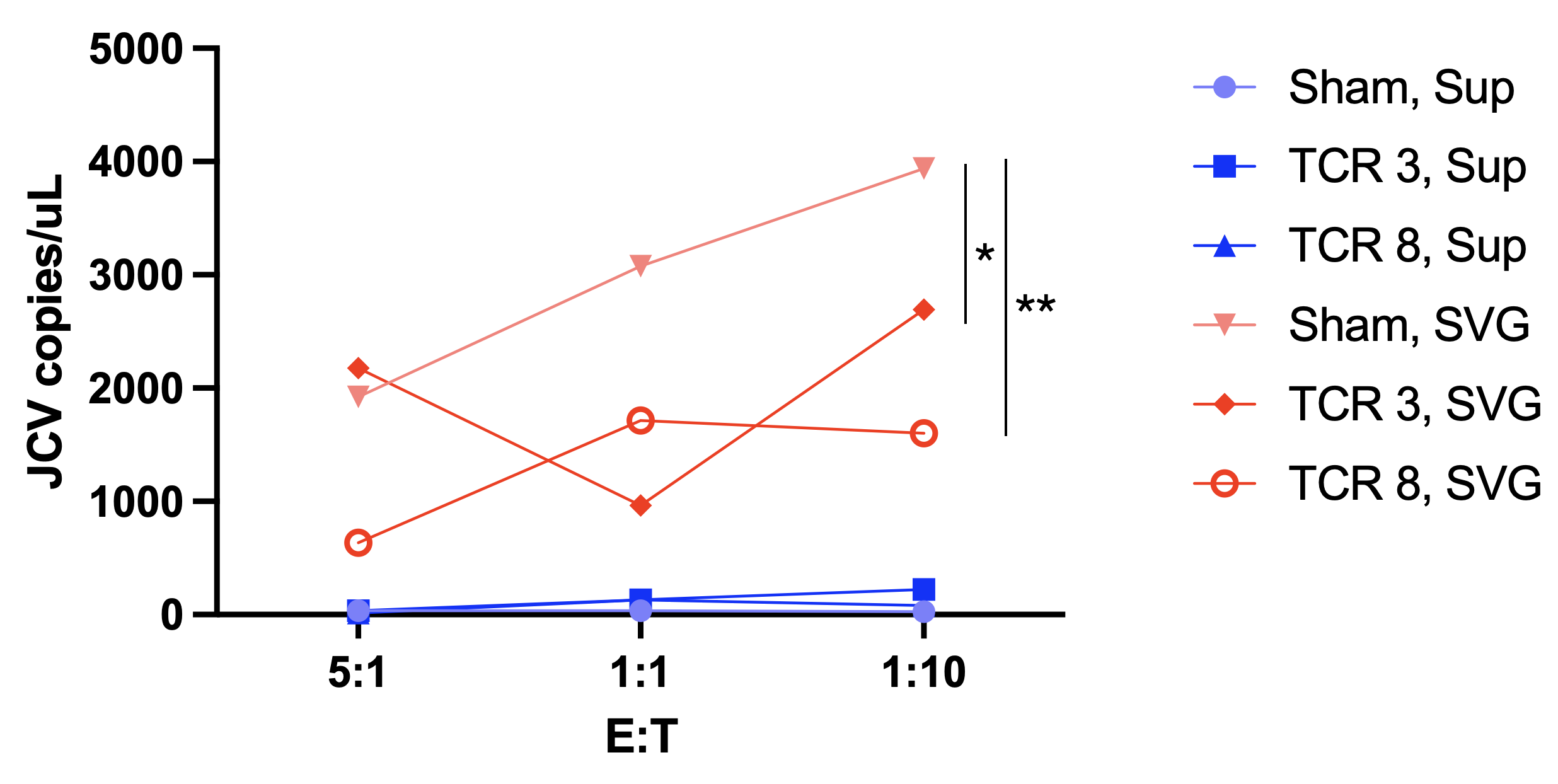
